# Feedback control of a two-component signaling system by an Fe-S-binding receiver domain

**DOI:** 10.1101/729053

**Authors:** Benjamin J. Stein, Aretha Fiebig, Sean Crosson

## Abstract

Two-component signaling systems (TCSs) function to detect environmental cues and transduce this information into a change in transcription. In its simplest form, TCS-dependent regulation of transcription entails phosphoryl-transfer from a sensory histidine kinase to its cognate DNA-binding receiver protein. However, in certain cases, auxiliary proteins may modulate TCSs in response to secondary environmental cues. *Caulobacter crescentus* FixT is one such auxiliary regulator. FixT is composed of a single receiver domain and functions as a feedback inhibitor of the FixL-FixJ (FixLJ) TCS, which regulates the transcription of genes involved in adaptation to microaerobiosis. We sought to define the impact of *fixT* on *Caulobacter* cell physiology and to understand the molecular mechanism by which FixT represses FixLJ signaling. *fixT* deletion results in excess production of porphyrins and premature entry into stationary phase, demonstrating the importance of feedback inhibition of the FixLJ signaling system. Although FixT is a receiver domain, it does not affect dephosphorylation of the oxygen-sensor kinase FixL or phosphoryltransfer from FixL to its cognate receiver FixJ. Rather, FixT represses FixLJ signaling by inhibiting the FixL autophosphorylation reaction. We have further identified a 4-cysteine motif in *Caulobacter* FixT that binds an Fe-S cluster and protects the protein from degradation by the Lon protease. Our data support a model in which oxidation of this Fe-S cluster promotes degradation of FixT *in vivo*. This proteolytic mechanism facilitates clearance the of the FixT feedback inhibitor from the cell under normoxia and resets the FixLJ system for a future microaerobic signaling event.

**Importance:** Two-component signal transduction systems (TCSs) are broadly conserved in the bacterial kingdom and generally contain two molecular components: a sensor histidine kinase and a receiver protein. Sensor histidine kinases alter their phosphorylation state in direct response to a physical or chemical cue, whereas receiver proteins “receive” the phosphoryl group from the kinase to regulate a change in cell physiology. We have discovered that a single-domain receiver protein, FixT, binds an Fe-S cluster and controls *Caulobacter* heme homeostasis though its function as a negative feedback regulator of the oxygen-sensor kinase, FixL. We provide evidence that the Fe-S cluster protects FixT from Lon-dependent proteolysis in the cell and endows FixT with the ability to function as a second, autonomous oxygen/redox sensor in the FixL-FixJ signaling pathway. This study introduces a novel mechanism of regulated TCS feedback control by an Fe-S-binding receiver domain.

## Introduction

Environmental sensing in bacteria is often mediated by two-component signaling systems (TCSs), consisting of a sensory histidine kinase and a response regulator (1). In canonical TCSs, the sensory histidine kinase autophosphorylates a single conserved histidine residue in response to an environmental cue. This phosphoryl group is then transferred from the histidine to a conserved aspartate residue on the receiver (REC) domain of its cognate response regulator. The phosphorylated response regulator elicits a cellular response, generally via an effector domain at its C-terminus (1). In many cases, this effector domain directly binds DNA and regulates transcription. However, a subclass of response regulators, known as the single-domain response regulators (SDRRs), consist of only a REC domain and induce a cellular response via protein-protein interactions (2-6). Although the general model of TCSs is relatively simple, these so-called two-component systems can in fact be part of complex phosphorelays involving multiple proteins, including sensory histidine kinases, histidine phosphotransfer proteins, and SDRRs (7-11).

FixL-FixJ (FixLJ) is an oxygen-responsive TCS that incorporates a regulatory SDRR, FixT, to control system output (12-14). FixL is a histidine kinase whose autophosphorylation activity is regulated by oxygen binding to a heme iron center on its sensory PAS (Per-Arnt-Sim) domain (15-17). FixL has higher kinase activity under microaerobic conditions, which results in increased phosphorylation of its cognate response regulator, FixJ. Phosphorylated FixJ induces a microaerobic-specific transcriptional program both directly via DNA binding and indirectly via upregulation of the transcription factor *fixK* (18, 19). FixL and FixJ were first identified in *Sinorhizobium meliloti* and have been most extensively characterized in diazotrophic rhizobiales (20, 21). In this group of bacteria, FixLJ primarily regulates microaerobic respiration and nitrogen fixation, a process that is extremely sensitive to oxygen levels. In addition, some rhizobiales employ FixLJ to regulate heme biosynthesis, anaerobic nitrate respiration, and expression of hydrogenases (22). In select organisms, FixLJ also upregulates the SDRR *fixT*, which functions as feedback inhibitor of the FixL-FixJ system (12, 13).

Initial biochemical work demonstrated that *S. meliloti* FixT directly inhibits accumulation of autophosphorylated FixL (14). This inhibition may arise via effects on the histidine autophosphorylation reaction and/or the reverse reaction with ADP. In addition, *in vitro* experiments did not detect FixT phosphorylation or inhibition of phosphotransfer from FixL to FixJ (14). Thus, despite having the primary structure of an SDRR, FixT does not apparently act as a competing receiver domain in *S. meliloti*. Interestingly, a subsequent genetic screen determined that *S. meliloti* FixT activity requires the presence of an additional gene, *asnO*, that is related to glutamine amidotransferase proteins and involved in N-acetylglutaminylglutamine synthesis (23, 24). This finding raises the possibility that FixT may sense metabolic signals and integrate information into the FixLJ circuit. However, the mechanism by which AsnO affects FixT activity is not understood.

*fixL, fixJ*, and *fixT* are prevalent in the order rhizobiales, but they have also been identified in *Caulobacter crescentus*, a free-living bacterium originally isolated from freshwater (12). *C. crescentus* is an obligate aerobe that does not fix nitrogen, and its FixLJ TCS primarily serves to activate expression of high affinity terminal oxidases and select metabolic pathways, including heme biosynthesis, under microaerobic conditions (12). Similar to *S. meliloti, C. crescentus* FixLJ strongly induces expression of the SDRR feedback inhibitor, *fixT* (21.5% sequence identity to *S. meliloti* FixT). In this study, we use the *C. crescentus* FixLJ system as a model to investigate the biological importance of TCS feedback control. Unlike *S. meliloti*, we observe distinct defects in a *C. crescentus* strain lacking *fixT*, highlighting the importance of negative feedback regulation of the FixLJ system. In addition, we conduct experiments aimed at defining the molecular mechanism by which FixT represses FixLJ signaling. Our results demonstrate that *C. crescentus* FixT preferentially inhibits the forward FixL autophosphorylation reaction, which is the biochemical basis of its function as an inhibitor of FixLJ-dependent transcription. Most notably, we identify a 4-cysteine (4-Cys) motif unique to *Caulobacter* FixT. This feature of primary structure supports binding of an Fe-S cluster and influences the function of FixT as an inhibitor of FixLJ. Mutational analysis implicates the Fe-S cluster in stabilizing FixT against degradation by the Lon protease, and we provide evidence that oxidation of the Fe-S cluster destabilizes FixT in the cell. We conclude that FixT is a novel Fe-S-binding SDRR capable of autonomously detecting changes in oxygen or the cellular redox state, and transducing those signals, via proteolytic susceptibility, into a change in FixLJ signaling.

## Results

### Loss of *fixT* affects porphyrin metabolism and growth

Feedback inhibition is a common mechanism to restrain biological signaling circuits (25, 26). Besides derepression of FixLJ-dependent transcription, no phenotypic consequences for loss of the *fixT* feedback inhibitor have been reported to date. We postulated that the physiological importance of FixT might be more evident in *C. crescentus* than in *S. meliloti*. Using a β-galactosidase transcriptional fusion assay, we found that the FixLJ system was active in cultures grown to stationary phase with limited aeration and, importantly, deletion of *fixT* resulted in strong derepression of FixLJ-dependent transcription (Fig. 1A). Under these same conditions Δ*fixT* cells exhibited a red pigmentation compared with wild-type cells (Fig. 1B). This color change was genetically complemented by ectopic *fixT* expression. To further characterize this color phenotype, we compared the absorbance spectra of lysates from wild-type and Δ*fixT* cells. These lysates exhibited a characteristic porphyrin absorbance spectrum, with a soret maximum at 414 nm and four *Q* bands at 506 nm, 542 nm, 580 nm, and 632 nm (Fig. 1C) (27). Importantly, the magnitude of these peaks was correlated with the degree of pigmentation observed in cell pellets (Fig 1B,C). Previous work found that the FixLJ TCS upregulates transcription of several genes involved in heme biosynthesis, and thus, it is likely that the porphyrin(s) we observe derive from this metabolic pathway (12). Moreover, the presence of 4 *Q* bands, rather than 2, indicates accumulation of free-base porphyrins, such as the heme precursor protoporphyrin IX (Fig. S1A) (27). In agreement with these data, excitation of wild-type and Δ*fixT* lysates with 404 nm light generated fluorescence maxima at 588 nm, 634 nm, and 698 nm (Fig. S1B). The latter two maxima suggest the presence of free-base porphyrin(s), whereas the peak at 588 nm may reflect porphyrin(s) bound to a closed-shell metal like Zn (28). This accumulation of free-base porphyrins in the Δ*fixT* strain suggests that heme biosynthesis may outpace iron availability. To test this hypothesis, we grew wild-type and Δ*fixT* cells in PYE media supplemented with 10 µM FeSO_4_. Lysates from these cells lost the fluorescence characteristics of free-base porphyrins (Fig. S1B). In addition, absorbance spectra of lysates from Fe-treated cells exhibited only 2 *Q* bands (Fig. S1C). These results provide evidence that, relative to wild-type cells, Δ*fixT* cells strongly upregulate heme biosynthesis and accumulate free-base porphyrin precursors due to insufficient iron availability.

**Figure 1:**
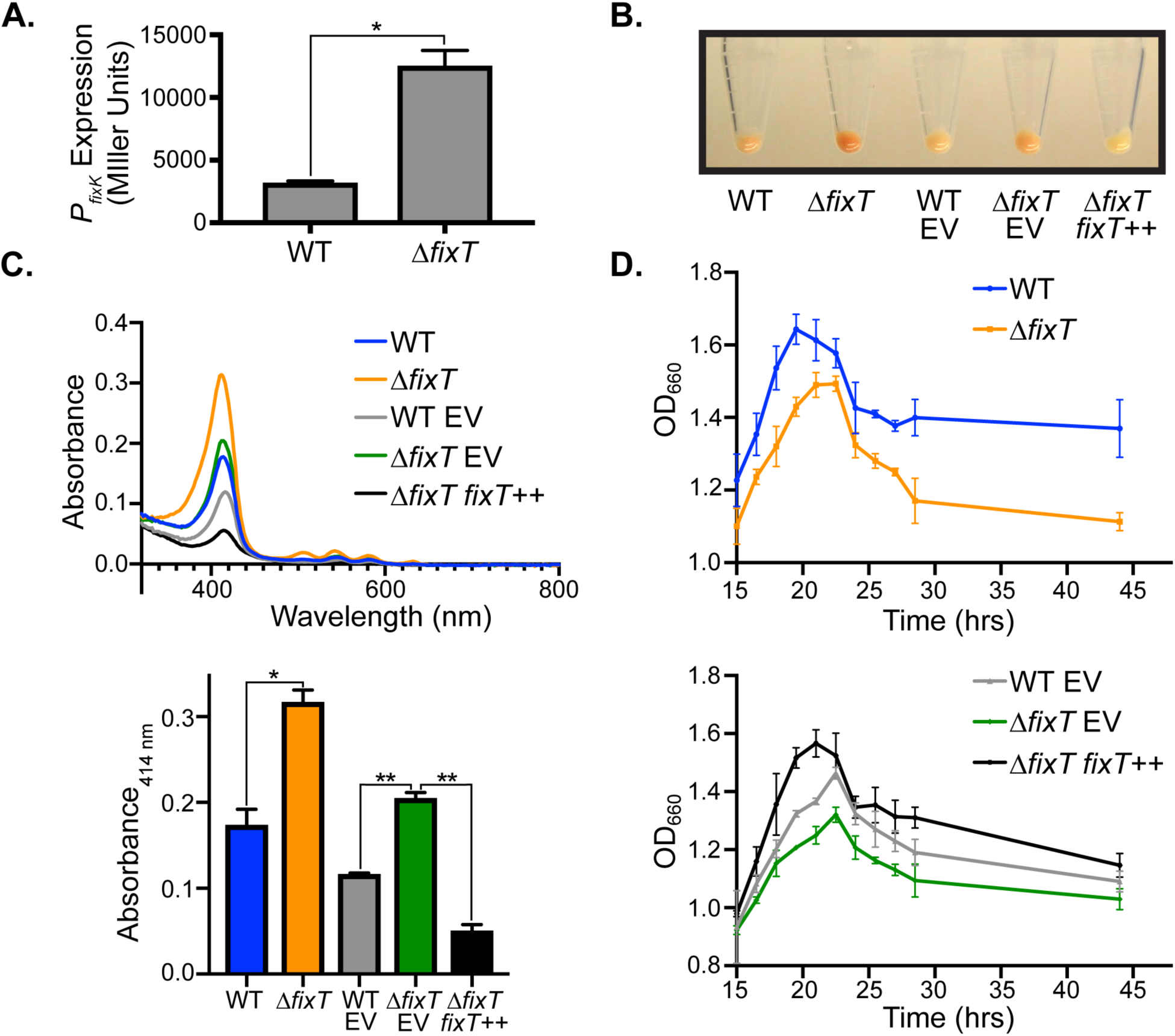
Loss of *fixT* leads to derepression of FixLJ-dependent transcription, protoporphyrin accumulation, and a stationary phase growth defect. (A) Activity of a *P*_*fixK*_-*lacZ* transcriptional reporter in wild-type (WT) or Δ*fixT* cells grown to high density with limited aeration (average ± SD, n = 3 biological replicates, each with 3 averaged technical replicates). * = *p* < 0.0005, Student t-Test. P_*fixK*_ is directly activated by FixLJ (12, 18, 19). (B) Cell pellets harvested from cultures grown as in A. Plasmid free wild-type (WT) and Δ*fixT* pellets are on the left. Strains carrying either empty pMT805 (EV) or xylose-inducible pMT805-*fixT* (*fixT*^++^) for complementation are on the right. (C) Top: Absorption spectra of cell lysates showing prominent soret peaks (414 nm) and 4 *Q* bands. Strain genotypes are annotated as in B. Bottom: Quantification of lysate absorbance at 414 nm (average ± SD, n = 3). * = *p* < 0.005, Student t-Test. ** = *p* < 0.0001, one-way ANOVA followed by Dunnett’s post-test comparison to Δ*fixT* EV. (D) Growth curve, measure by optical density (OD_660_), following entry into stationary phase for wild-type (WT) and Δ*fixT* cells (Top) and plasmid-bearing complementation strains (Bottom). Points are averages of 3 biological replicates ± SD.

In addition to exhibiting a color change consistent with the accumulation of heme precursors, Δ*fixT* cultures also displayed growth defects at high culture density. Specifically, the terminal optical density of Δ*fixT* cultures was lower than that of wild-type when grown with limited aeration (Fig. 1D). We observed this difference despite the fact that the exponential growth rate of wild-type and Δ*fixT* cultures was indistinguishable (Fig. S2). Ectopic overexpression of *fixT* complemented this stationary phase defect (Fig. 1D). We surmised that the inability of Δ*fixT* cells to grow to high density might be due to the accumulation of heme precursors observed in these cells, as high levels of free porphyrins are generally toxic (29). Alternatively, upregulation of heme biosynthesis might indirectly lead to a growth defect at high cell density by contributing to iron starvation. In agreement with either of these hypotheses, supplementation of iron in the growth media largely mitigated the high density growth defect in Δ*fixT* cells (Fig. S1D). We conclude that under microaerobic conditions with limited iron, loss of *fixT* leads to a detrimental accumulation of porphyrins in *C. crescentus*.

### FixT acts as a direct inhibitor of the FixL kinase

Biochemical studies of *S. meliloti* FixT indicated that it inhibits accumulation of autophosphorylated FixL (14). To examine whether *C. crescentus* FixT functions in a similar manner, we tested its inhibitory activity *in vitro* using purified FixT, soluble FixL (FixL_118-495_, missing the transmembrane domain), and FixJ(D55A) (a non-phosphorylatable mutant). Previous work noted that FixL autokinase activity is enhanced by FixJ, indicating that FixL and FixJ likely work as a complex *in vivo* (30, 31). Using [*γ*-^32^P]ATP, we observed that FixL robustly autophosphorylates over a 30-minute time course in the presence of FixJ(D55A). However, the addition of excess FixT inhibited accumulation of autophosphorylated FixL by ∼3-fold (Fig. 2A). This inhibitory activity did not require FixL stimulation by FixJ(D55A), indicating that FixT does not simply compete with FixJ(D55A) for binding to FixL (Fig. 2B). Moreover, FixT completely blocked activation of FixL by FixJ(D55A), even at equimolar excess FixJ(D55A) and FixT. Given the low micromolar apparent affinities of FixL for both FixT and FixJ(D55A), these data suggest that the biochemical effects of FixT are dominant to those of FixJ (Fig. 2B, 4B). Thus, similar to *S. meliloti* FixT, *C. crescentus* FixT inhibits accumulation of phosphorylated FixL in the presence or absence of FixJ. This inhibition may arise from effects on FixL autophosphorylation and/or FixL dephosphorylation.

**Figure 2:**
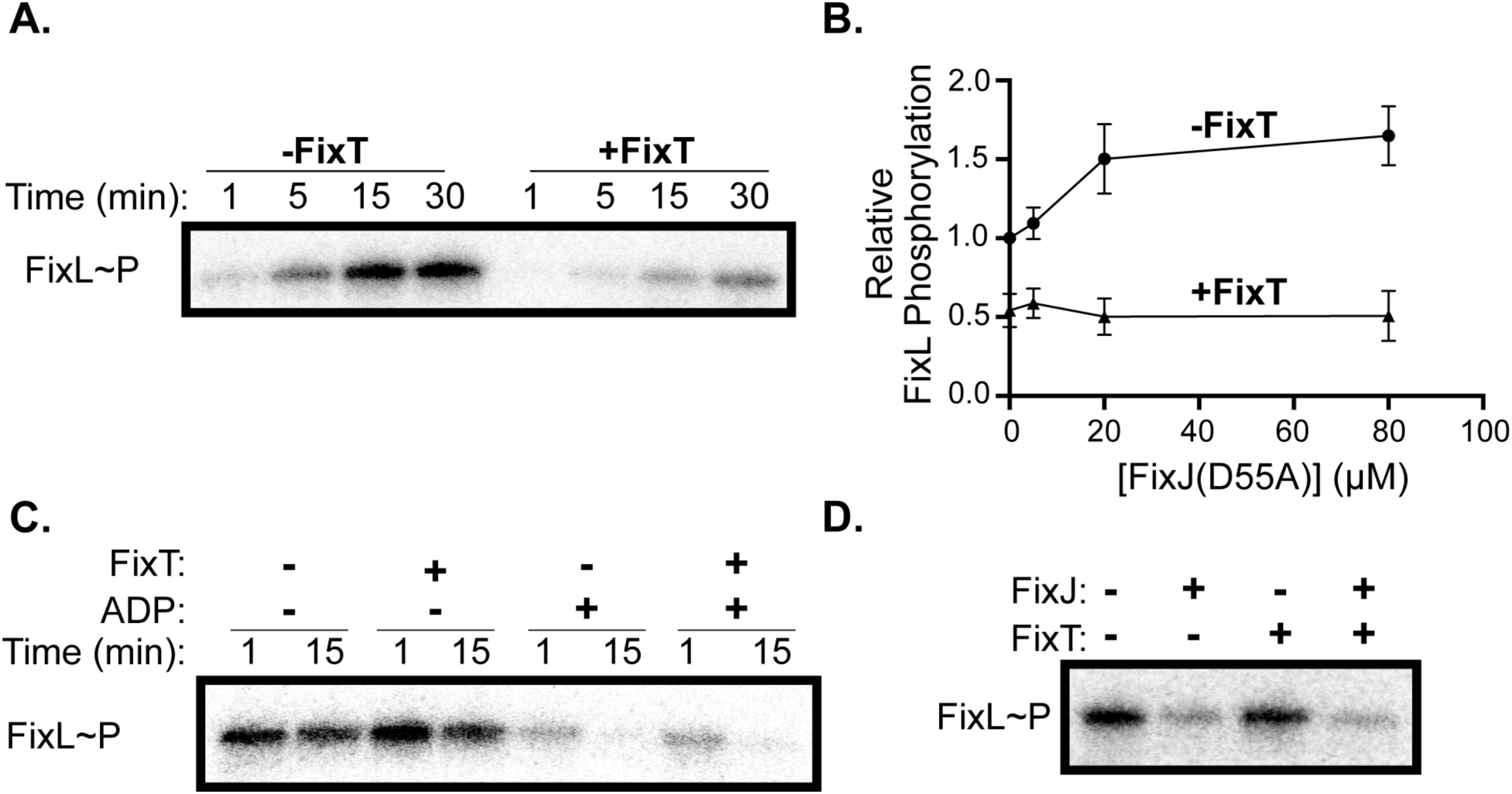
FixT inhibits FixL autophosphorylation *in vitro*. (A) Autophosphorylation of FixL (5 µM) in the presence or absence of FixT (80 µM). Reactions were carried out in the presence of FixJ(D55A) (20 µM). (B) FixL (5 µM) autophosphorylation with increasing FixJ(D55A) concentrations in the presence or absence of FixT (80 µM). Reactions were quenched after 15 min, separated by SDS-PAGE, and phosphorylated-FixL was quantified by phosphorimaging. Values were normalized to the reaction without FixT and FixJ, average ± SD, n = 3. (C) Disappearance of phosphorylated FixL (5 µM), purified from nucleotide, in the presence of excess FixT (80 µM) and/or ADP (0.5 mM). (D) Disappearance of phosphorylated FixL (5 µM) 30 seconds after addition of FixJ (2.5 µM) in the presence or absence of excess FixT (80 µM).

**Figure 3:**
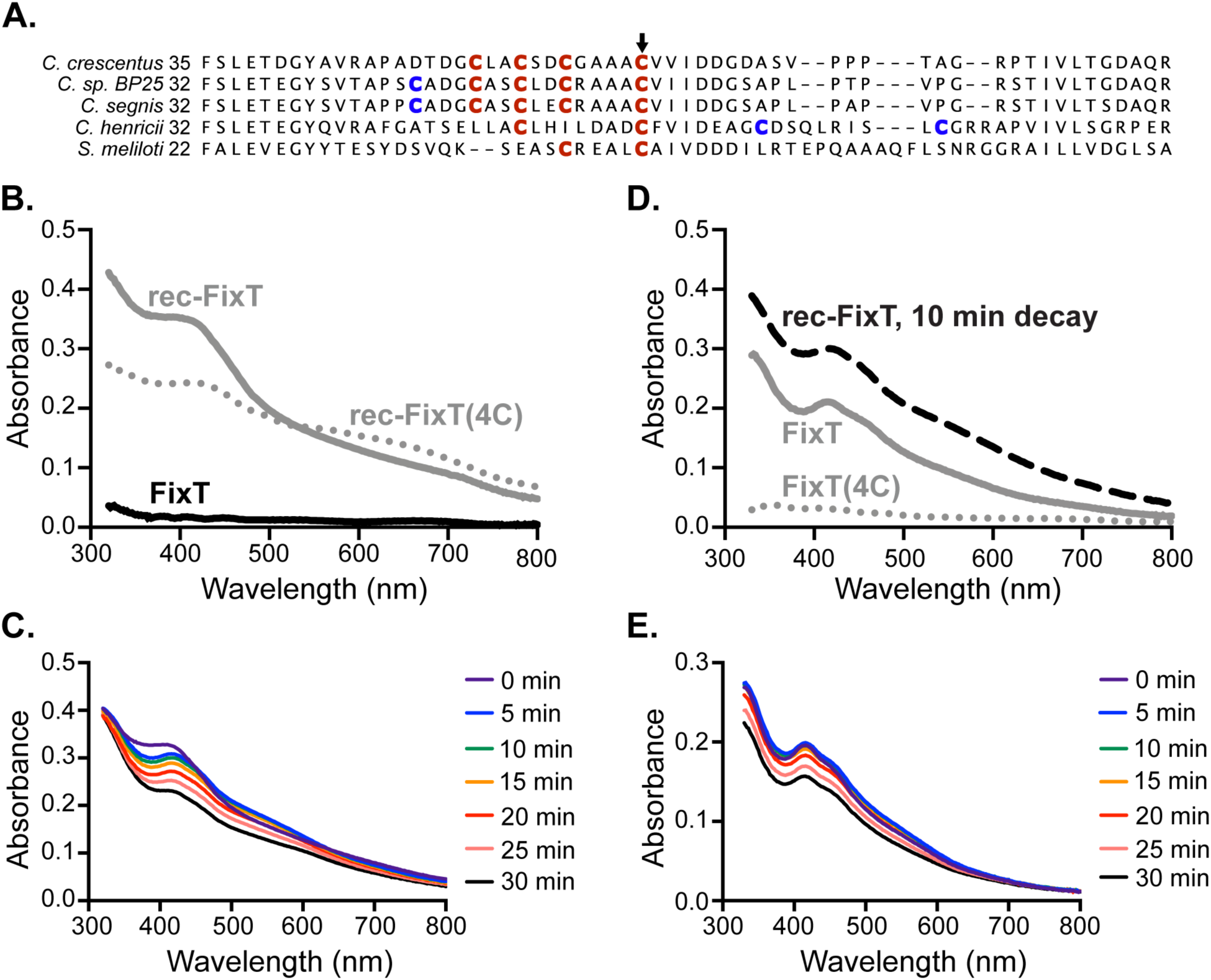
FixT binds an Fe-S cluster. (A) Sequence alignment of FixT proteins from several *Caulobacter* species and *S. meliloti*. The position of the first amino acid residue of each sequence is on the left. Cys residues aligning with the *C. crescentus* 4-Cys motif are in red. Additional nearby Cys residues are in blue. The position corresponding to *C. crescentus* C64 is marked by an arrowhead. (B) Absorption spectra of aerobically-purified FixT (FixT), anaerobically-reconstituted wild-type FixT (rec-FixT), and anaerobically-reconstituted FixT(4C) (rec-FixT(4C)). (C) Absorption spectra of rec-FixT collected at 5 minute intervals after exposure to ambient air. (D) Absorption spectra of anaerobically-purified H_6_-Sumo-FixT and H_6_-Sumo-FixT(4C) at 400 µM. The 10 min decay curve of FixT with a reconstituted Fe-S cluster (rec-FixT) from panel C is shown for comparison. (E) Absorption spectra of anaerobically-purified H_6_-Sumo-FixT collected at 5 minute intervals after exposure to ambient air.

**Figure 4:**
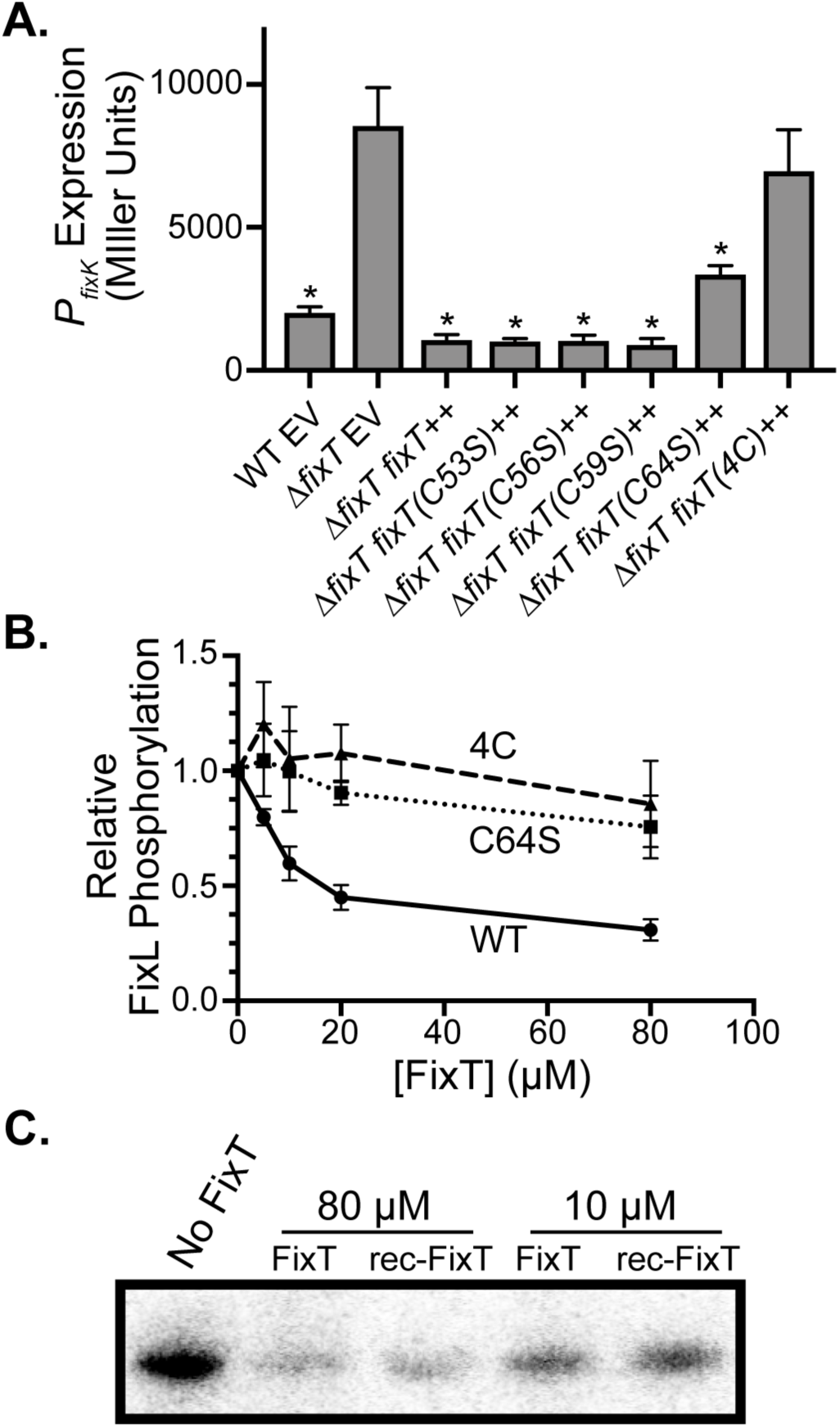
The 4-Cys motif, but not the Fe-S cluster, is necessary for FixL inhibition *in vitro*. (A) *P*_*fixK*_*-lacZ* transcriptional reporter activity in strains carrying an empty vector (EV) or a plasmid overexpressing wild-type or mutant *fixT* alleles (average ± SD, n = 3 biological replicates, each with 3 averaged technical replicates). * = *p* < 0.0001, one-way ANOVA followed by Dunnett’s post-test comparison to Δ*fixT* EV. (B) Relative FixL (5 µM) autophosphorylation, in the presence of FixJ(D55A) (20 µM), with increasing concentrations of wild-type (WT) or mutant FixT. Phosphorylation was measured as in Fig 2B and normalized to the reaction without FixT, average ± SD, n = 3. (C) Autophosphorylation of FixL (5 µM) in the presence of FixT or rec-FixT. Phosphorylation reactions were performed under anaerobic conditions.

To address the effect of FixT on FixL dephosphorylation (either hydrolysis or a reverse reaction with ADP), we examined the disappearance of phosphorylated FixL after removing nucleotides by desalting. Addition of excess FixT had no effect on the slow loss of FixL-phosphate observed over 15 minutes, indicating that FixT does not act as a phosphatase (Fig. 2C). Addition of excess ADP led to rapid loss of the phosphoryl group from FixL, perhaps by a reverse reaction to form ATP (32). The presence of excess FixT did not affect this phosphoryl loss (Fig. 2C). Taken together, these results indicate that FixT primarily reduces accumulation of phosphorylated FixL by inhibiting the FixL autophosphorylation reaction, rather than stimulating loss of FixL-phosphate.

We also examined whether FixT could interfere with phosphoryltransfer from FixL to FixJ. To test this possibility, we monitored dephosphorylation of FixL after the addition of FixJ. Addition of FixJ alone induced a rapid (within 30 seconds) loss of FixL-phosphate. Moreover, a 32-fold molar excess of FixT did not inhibit dephosphorylation of FixL by FixJ (Fig. 2D). Thus, in agreement with our results with FixJ(D55A), these data indicate that FixT does not interfere with the interaction between FixL and FixJ. This observation, paired with the fact that we never observed phosphorylation of FixT in our experiments (data not shown), provides evidence that FixT does not act as a competitive phosphoryl sink.

### FixT binds an Fe-S cluster *in vitro*

Our biochemical dissection of FixT activity indicated that it functions similarly to *S. meliloti* FixT (14). However, we observed that *C. crescentus* FixT, unlike *S. meliloti* FixT, contains a 4-Cys motif (C53, C56, C59, C64) reminiscent of iron-sulfur (Fe-S) binding motifs in other proteins (Fig. 3A) (33, 34). We also noted that *C. crescentus* FixT was light brown at early stages of purification, indicating the potential presence of bound iron. Given these observations, we hypothesized that FixT may bind an Fe-S cluster. To test this possibility, we attempted to chemically reconstitute an Fe-S cluster on aerobically purified FixT in an anaerobic chamber. Upon reconstitution, FixT adopted a dark brown color reminiscent of many Fe-S proteins (35-37). Absorption spectra of this reconstituted sample (rec-FixT) exhibited a shoulder in the blue region of the visible spectrum (*λ* ∼ 420 nm), similar to previously reported spectra for [4Fe-4S] cluster-binding proteins (Fig. 3B) (33). Importantly, these spectral features were not evident in non-reconstituted FixT (Fig. 3B). Fe measurements using ferrozine detected an average of 3.8 ± 0.3 Fe atoms per rec-FixT monomer (average ± SD, n = 3). These data indicate that FixT is capable of binding an Fe-S cluster, most likely in the form of a [4Fe-4S] cluster. Additionally, we mutated each position in the 4-Cys motif to Ser (FixT(4C)) and tested the ability of this mutant protein to bind a reconstituted Fe-S cluster. Reconstitution reactions with FixT(4C) produced granular black material indicating formation of iron sulfides instead of Fe-S clusters. Rec-FixT(4C) also exhibited relatively strong absorbance beyond 650 nm, further evidence of significant iron-sulfide formation (Fig. 3B) (33). We conclude that the FixT 4-Cys motif is important for productive reconstitution of an Fe-S cluster.

We surmised that the FixT Fe-S cluster is likely sensitive to oxygen exposure as *a)* we observe little pigmentation and absorbance in aerobically purified FixT and *b)* most solvent-accessible [4Fe-4S] clusters react with molecular oxygen (33, 38, 39). To test his hypothesis, we measured absorption spectra after exposing rec-FixT to the ambient atmosphere. We observed a spectral transition within the first 5 minutes characterized by a red-shift of the major absorbance peak, a shoulder at 465 nm, and increased absorbance at ∼540 nm. This initial transition was followed by a slow overall loss in absorbance (Fig. 3C). These changes are consistent with decay to a [2Fe-2S] cluster, followed by a slower loss of remaining iron and sulfide (40-42). Thus, like many Fe-S clusters, the FixT cluster is sensitive to atmospheric concentrations of molecular oxygen, raising the possibility that it acts as a sensor of cytoplasmic oxygen levels.

To examine whether FixT can acquire this oxygen-sensitive Fe-S cluster in cells, we anaerobically purified FixT and FixT(4C) (retaining their H_6_-Sumo tags), heterologously expressed in *E. coli*, and examined their absorption spectra. The absorption spectrum of anaerobically-purified FixT was most similar to the decay spectra for rec-FixT, with an absorption peak at 420 nm and a shoulder at 465 nm (Fig. 3D). In contrast, anaerobically-purified FixT(4C) exhibited no significant absorbance at 420 nm, indicating that the 4-Cys motif is indeed required for Fe-S cluster binding. Consistent with these spectra, we detected Fe in the FixT sample (0.21 ± 0.07 Fe atoms per FixT monomer, n = 2), but not in the FixT(4C) sample (−0.01 ± 0.02 Fe atoms per FixT(4C) monomer, n = 2). Similar to rec-FixT, exposure to ambient atmosphere led to cluster decay in anaerobically-purified FixT, with an early transition marked by an increase in absorbance at ∼540 nm (Fig. 3E). Thus, a subfraction of FixT purified from *E. coli* contains an Fe-S cluster similar to that seen in our *in vitro* reconstitution experiments.

### The FixT 4-Cys motif plays an important role in feedback inhibition

We next sought to test the importance of the 4-Cys motif for the regulatory function of FixT *in vivo*. Using the *P*_*fixK*_*-lacZ* fusion (a reporter of FixLJ-dependent transcription) and microaerobic growth conditions, we measured the effect of individual and combined Cys-to-Ser mutations on the ability of FixT to repress FixLJ output. Ectopic overexpression of *fixT* strongly repressed transcription from the *P*_*fixK*_*-lacZ* reporter plasmid (Fig. 4A). Overexpression of FixT C53S, C56S, and C59S single mutants repressed transcription similarly to wild-type FixT. However, the C64S single mutant only partially repressed *P*_*fixK*_*-lacZ* activity. We note that this cysteine is broadly conserved in FixT proteins, unlike other residues in the 4-Cys motif (Fig. 3A). Importantly, mutation of all four Cys residues (FixT(4C)) abolished repression entirely (Fig. 4A). Thus, the 4-Cys motif is critical for FixT function *in vivo*.

To examine the importance of these Cys residues for function *in vitro*, we purified the C64S and 4C mutant proteins and tested their inhibitory activities against FixL. These FixT mutants were both defective at inhibiting FixL autophosphorylation, even at 32-fold molar excess (Fig. 4B). At these concentrations, we did not observe significant differences between the mutant proteins, as might be expected from the *in vivo* β-galactosidase reporter assay (Fig. 4A). It is possible that the dynamic range of our *in vitro* assay is not adequate to robustly detect modest inhibition by the C64S mutant protein. Nevertheless, our data clearly demonstrate that the 4-Cys motif, and C64 in particular, is critical for FixT inhibitory activity.

Given that the 4-Cys motif is important for forming an Fe-S cluster, we next tested whether this Fe-S cluster might affect the ability of FixT to inhibit FixL autophosphorylation *in vitro*. Interestingly, aerobically-purified FixT (FixT) and Fe-S-bound FixT (rec-FixT) similarly inhibited FixL autophosphorylation under anaerobic conditions (Fig. 4C). Thus, FixT does not require the Fe-S cluster to inhibit FixL autokinase activity *in vitro*.

### FixT protein levels are regulated by Lon proteolysis

Although the Fe-S cluster *per se* does not affect the activity of FixT as a FixL inhibitor, it may play an important structural role. In fact, many Fe-S clusters affect protein folding, thermodynamic stability, and/or proteolytic stability (43, 44). We also hypothesized that proteolysis may play a significant role in regulating FixT as a feedback inhibitor, because FixT should be cleared from the cell under aerobic conditions to allow future microaerobic transcriptional responses. To probe FixT proteolytic stability, we first examined steady-state FixT levels in strains carrying deletions of energy-dependent protease genes. To avoid complications that might arise from endogenous transcriptional regulation of *fixT*, we deleted the native *fixT* gene and monitored FixT expressed from a xylose-inducible promoter. Western blots of gel-resolved lysates from these strains detected low levels of FixT in all but Δ*fixT*Δ*lon*, where FixT levels were dramatically higher (Fig. 5A). We conclude that the Lon protease plays an important role in the degradation of FixT in *C. crescentus*. To ensure that this result was not an artifact of *fixT* overexpression, we also examined natively-expressed FixT levels in WT and Δ*lon* cultures grown to saturation. Natively-expressed FixT was below the limit of detection in wild-type cells, but observable in Δ*lon* cells (Fig. 5B). As a control, we also demonstrated that FixT signal was absent from Δ*fixT*Δ*lon* cells. Together these results indicate that Lon regulates FixT steady-state levels *in vivo*. To more directly assess proteolysis by Lon, we examined the half-life of FixT, in *lon*^*+*^ and Δ*lon* backgrounds, after inhibiting translation with oxytetracycline. Due to low protein levels at stationary phase (presumably as a result of reduced translation rates), we conducted these experiments using mid-exponential phase cells. Upon addition of oxytetracycline, we observed loss of FixT in the *lon*^*+*^ background with a half-life of 42 ± 4.9 min (Fig. 5C). Importantly, loss of FixT was slower in Δ*lon* cells (half-life not determined, but > 60 min). We also assayed the *in vivo* stability of FixT(4C) and found that it was degraded more rapidly than the wild-type protein, with a half-life of 13 ± 3.9 min (Fig. 5C). As expected, this degradation was also *lon*-dependent. Taken together, our results support a model in which FixT is degraded by Lon *in vivo* and that loss of the 4-Cys motif enhances degradation. Given our *in vitro* reconstitution experiments, these data also suggest that the Fe-S cluster may protect FixT against Lon proteolysis. To test this hypothesis, we measured the half-lives of FixT and FixT(4C) in cultures with enhanced aeration, as we have shown that the FixT Fe-S cluster is sensitive to oxygen *in vitro*. To enhance aeration, we reduced the culture volume from 10 mL to 4 mL to increase head space and increased the tilt angle during shaking. FixT was degraded more quickly under these conditions, with a ∼2-fold shorter half-life (24 min vs 42 min, p < 0.05, Student t test). In contrast, FixT(4C) degradation was not significantly affected by increased aeration. Thus, FixT proteolytic stability is sensitive to the degree of culture aeration if the 4-Cys motif is present.

**Figure 5:**
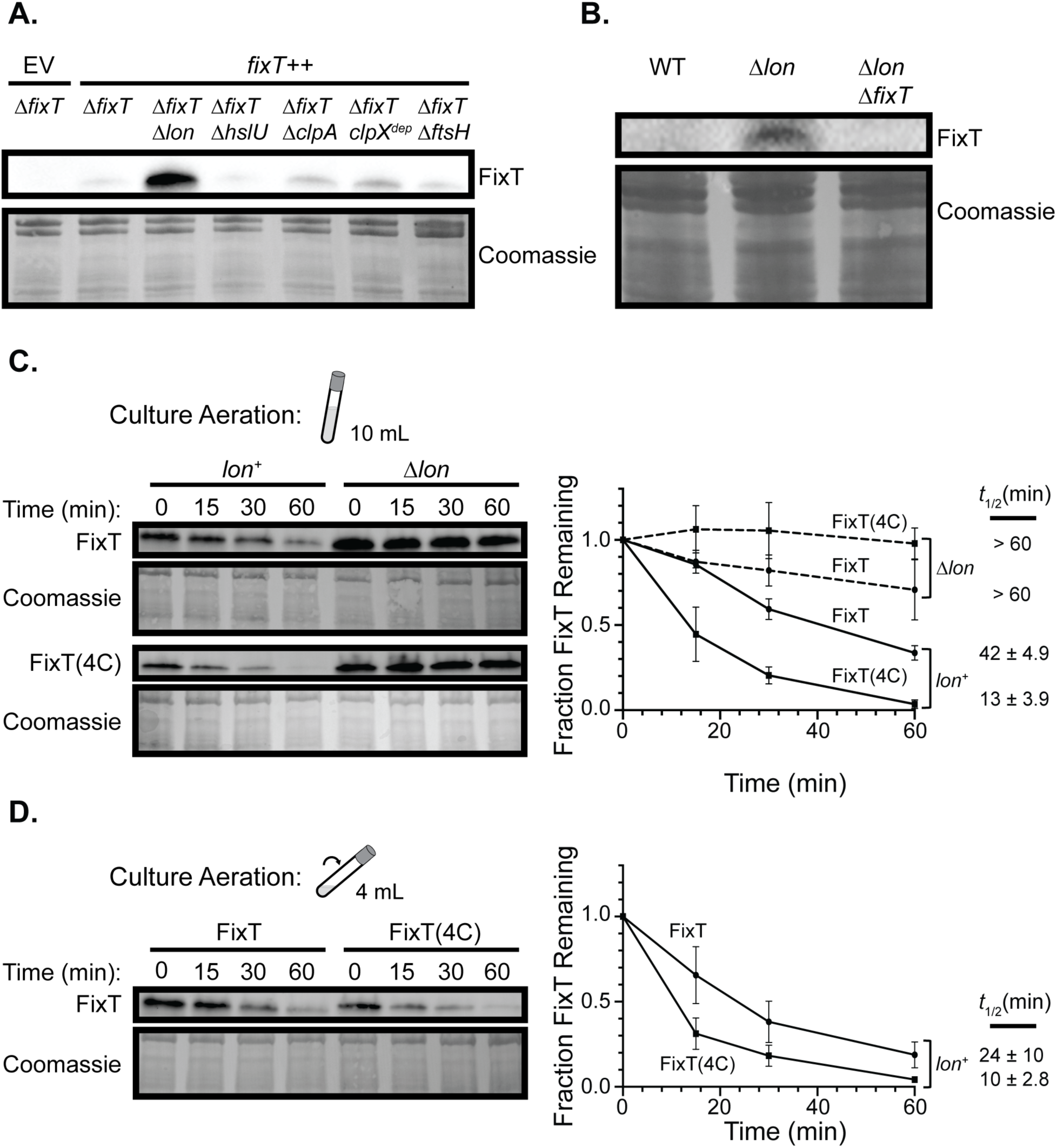
The Lon protease degrades FixT *in vivo*. (A) Western blot of lysates from strains lacking ATP-dependent protease systems probed with anti-FixT antibody. Strains lack chromosomal *fixT* and carry either an empty vector (EV) or a *fixT* overexpression plasmid (*fixT*^++^). Coomassie stain of the membrane is shown as a loading control. (B) Western blot using an anti-FixT antibody to assess steady-state levels of endogenously-expressed *fixT* in wild-type (WT), Δ*lon*, and Δ*fixT*Δ*lon* cell lysates under saturating, microaerobic conditions. Coomassie stain of the membrane is shown as a loading control. (C) *In vivo* degradation experiments using exponential phase cells grown under limited aeration (determined by culture volume and tilt depicted in cartoon). Left: Representative anti-FixT western blots of Δ*fixT* (*lon*^+^) and Δ*fixT*Δ*lon* (Δ*lon*) cells overexpressing *fixT* or *fixT(4C)*. Cells were sampled at the indicated intervals after arresting translation with oxytetracycline. Coomassie stains of membranes are shown as a loading control. Right: Quantification of FixT levels from replicates of the experiment shown to the left (average ± SD, n = 3). The half-life (*t*_1/2_, average ± SD, n = 3) of FixT in each strain is displayed to the right of the graph. The difference between FixT and FixT(4C) half-lives in the *lon*^+^ background is significant (*p* < 0.005, Student t-Test). (D) *In vivo* degradation experiments as in panel C using exponential phase cells grown under enhanced aeration (smaller volume and greater tilt angle as shown in cartoon). Left: Representative anti-FixT western blot of Δ*fixT* (*lon*^+^) cells overexpressing either *fixT* or *fixT(4C)*. Right: Quantification of FixT levels from replicates of the experiment shown to the left. Points and half-lives are averages ± SD, n = 3.

## Discussion

### The importance of feedback inhibition in the FixLJ system

Deletion of *fixT* in *C. crescentus* leads to the accumulation of porphyrins and premature entry into stationary phase (Fig. 1, S1). These are the first reported phenotypes for loss of FixT-dependent feedback inhibition in any organism. These phenotypes are also apparently linked, as supplementation of the growth medium with iron alters the absorbance of accumulated porphyrins and alleviates the growth defect. Why are Δ*fixT* phenotypes apparent in *C. crescentus* while none have been reported in *S. meliloti?* Previous studies found that FixLJ upregulates *hemA, hemB, hemE*, and *hemN* in *C. crescentus* (12). In contrast, *S. meliloti* FixLJ only significantly upregulates *hemN* (45). We suggest that multi-level regulation of heme biosynthesis by *C. crescentus* FixLJ underlies the porphyrin accumulation defect. This link between unchecked FixLJ-dependent transcription and porphyrin accumulation highlights an important physiological role for FixT in *C. crescentus*. Over-induction of heme biosynthesis is likely metabolically taxing, particularly in oligotrophic environments where iron levels may be limiting. Moreover, in low-iron environments, heme over-induction leads to accumulation of potentially toxic porphyrin intermediates (29). Thus, in *C. crescentus*, FixT may function largely to restrain activation of heme biosynthesis by FixLJ, balancing heme production with the levels of available nutrients, including iron.

### On the mechanism of FixT inhibition

Although the physiologic importance of FixT is more readily apparent in *C. crescentus* than in *S. meliloti*, its mechanism of action is similar in both organisms. Our results provide evidence that *C. crescentus* FixT primarily inhibits the FixL autophosphorylation reaction, as we observed no effect of FixT on FixL-phosphate hydrolysis or the reverse reaction with ADP (Fig. 2). FixT may inhibit FixL autophosphorylation by interfering with ATP binding. However, our inhibition experiments were conducted using a large excess of ATP (1 mM), making this model unlikely. Moreover, FixT does not appear to interfere with the binding of ADP to FixL. Therefore, the data are most consistent with a model in which FixT affects the catalytic activity of the FixL kinase domain.

These observations are largely in agreement with those for *S. meliloti* FixT. Although Garnerone et al. did not examine the reverse reaction with ADP, they observed that FixT inhibits accumulation of autophosphorylated FixL (14). We note that the degree of inhibition observed in their experiments was greater than in ours. This discrepancy may reflect inherent differences in inhibitory activity or FixL affinity between the two homologous systems, and thus evolutionary tuning of the degree of feedback in different organisms. Alternatively, this discrepancy may simply be a result of technical differences in the assay conditions or the particular constructs used in each *in vitro* system.

Our characterization of FixT activity places it in contrast to other well-studied histidine kinase inhibitors, such as KipI and Sda, which block both autophosphorylation and the reverse reaction with ADP (46, 47). These inhibitors appear to sterically interfere with communication between the ATP binding site on the catalytic domain and the histidine phosphorylation site, and thus, affect the phosphorylation reaction in both directions (47, 48). In contrast, our data suggest that FixT preferentially inhibits the forward autophosphorylation reaction. Perhaps FixT preferentially binds to and stabilizes unphosphorylated FixL, thereby largely affecting only the autophosphorylation reaction. Further biochemical and structural studies are necessary to characterize the interaction between FixL and FixT and reveal the molecular basis of inhibition.

### The FixT 4-Cys motif, Fe-S cluster, and proteolysis

Despite sharing a common mechanism of kinase inhibition, *C. crescentus* and *S. meliloti* FixT differ in the presence of a 4-Cys motif. In fact, the 4-Cys motif is largely conserved within the *Caulobacter* genus, but is absent in non-*Caulobacter* FixT proteins (Fig. 3A). Our *in vivo* and *in vitro* data provide evidence that this 4-Cys motif is important for both Fe-S cluster binding and FixT function as a FixL inhibitor (Fig. 3, 4). To our knowledge, *C. crescentus* FixT represents the first REC domain found to bind an Fe-S cluster. Thus, FixT may be added to a growing list of SDRR proteins with noncanonical ligands and/or modifications (6, 49, 50). The stoichiometry of this Fe-S cluster remains an open question; spectroscopic techniques such as EPR or Mössbauer spectroscopy are necessary to definitively determine cluster type (41). However, our initial spectroscopic data and Fe measurements are consistent with FixT binding a [4Fe-4S] cluster.

Although our data indicate that FixT is capable of binding an Fe-S cluster, we did not detect a significant difference in the ability of FixT and Fe-S-reconstituted FixT (rec-FixT) to inhibit FixL autophosphorylation *in vitro*. The 4-Cys mutant deficient in Fe-S cluster binding (FixT(4C)) is also defective in FixL inhibition, thus we cannot discern whether the Fe-S cluster *per se* contributes to inhibitory activity *in vivo*. It is possible that our Fe-S-reconstituted FixT does not fully reflect the form found *in vivo*, and therefore does not capture relevant effects of the Fe-S cluster on inhibitory activity. That said, our data do provide evidence that the Fe-S cluster plays a significant role in controlling FixT degradation by the Lon protease.

Lon is most widely known for its role in degrading damaged, unfolded, and misfolded proteins in the cytoplasm (51, 52). However, Lon is also known to degrade a variety of specific, folded substrates including SulA and DnaA (53-55). Our work demonstrates that FixT is also a substrate of Lon *in vivo*. As yet, we do not know the signal(s) that target FixT for degradation. Lon may recognize folded FixT via specific degron(s) or may simply degrade FixT upon full or partial unfolding. We can infer, however, that this degradation is likely to be physiologically relevant, as tuning FixT steady-state levels by deletion or overexpression has observable effects on FixLJ-dependent transcription (Fig. 1, 4). Thus, FixT levels are positioned such that changes in *fixT* expression and/or the rate of Lon degradation will affect FixLJ signaling.

Notably, the assembly states of Fe-S ligands have been shown to influence degradation rates of several proteins (44, 56, 57). For example, *E. coli* FNR is a [4Fe-4S]-binding transcription factor that is active under anaerobic conditions. Upon oxygen exposure, the FNR cluster degrades, leading to monomerization and degradation by the ClpXP protease (40, 44). We propose that the FixT Fe-S cluster may play a similar role in impeding degradation by the Lon protease. Our *in vivo* results clearly show that the FixT(4C) mutant, which does not efficiently reconstitute an Fe-S cluster *in vitro*, is degraded 3-4-fold faster than the wild-type protein (Fig. 5). Moreover, enhancing culture aeration, and thus presumably dissolved oxygen levels, reduces the half-life of wild-type FixT. Given the oxygen sensitivity of the FixT Fe-S cluster, these results strongly suggest that Fe-S-bound FixT resists Lon proteolysis. We hypothesize that the Fe-S cluster may increase FixT thermodynamic stability or induce conformational or oligomeric changes that block degron recognition.

Mounting evidence in bacteria and eukaryotes points to an ancient role for Lon in degrading Fe-S proteins and regulating Fe-S assembly pathways. In *Salmonella enterica*, Lon selectively degrades the Fe-S protein FeoC under aerobic conditions, thereby destabilizing FeoB and reducing ferrous iron transport (56). A recent proteomic study in *E. coli* identified several putative Lon substrates that bind Fe-S cofactors, as well as the Fe-S assembly protein IscS (58). In eukaryotes, multiple substrates of the mitochondrial Lon homologue have been identified, many of which are Fe-S binding metabolic enzymes or Fe-S cluster assembly proteins (57, 59-64). Moreover, disrupting mitochondrial Fe-S clusters, either genetically or via oxidative stress, enhances degradation of several Fe-S-binding Lon substrates (57, 59). Lon-family proteases may be particularly suited to detecting damage in Fe-S-binding proteins, as loss of Fe-S clusters can induce conformational changes that reveal hydrophobic regions, which are canonical Lon recognition sites (43, 65).

Importantly, regulation of FixT levels by proteolysis provides an additional node for signal integration into the FixLJ TCS (Fig. 6). We expect that the FixT Fe-S cluster will respond to physiochemical changes in the cytoplasm. We have shown that the FixT Fe-S cluster is sensitive to ambient air, and thus perhaps oxygen (or other oxidizing agents) triggers rapid FixT degradation, allowing reset of the FixLJ system upon return to aerobic growth. Moreover, differing affinities of FixL and the FixT Fe-S cluster for oxygen could significantly alter the “on” and “off” kinetics of the FixLJ system. Yet, Fe-S clusters are also known to respond to other inputs, such as nitric oxide stress and the status of iron-sulfur assembly pathways (which can, in turn, reflect iron availability or oxidative stress) (39). Further work is required to more comprehensively determine the signals sensed *in vivo* by the FixT Fe-S cluster. Finally, Lon itself may relay environmental information into the FixLJ system, as factors like cell cycle stage, adaptor proteins, and/or substrate competition might affect its activity towards FixT (68, 69).

**Figure 6:**
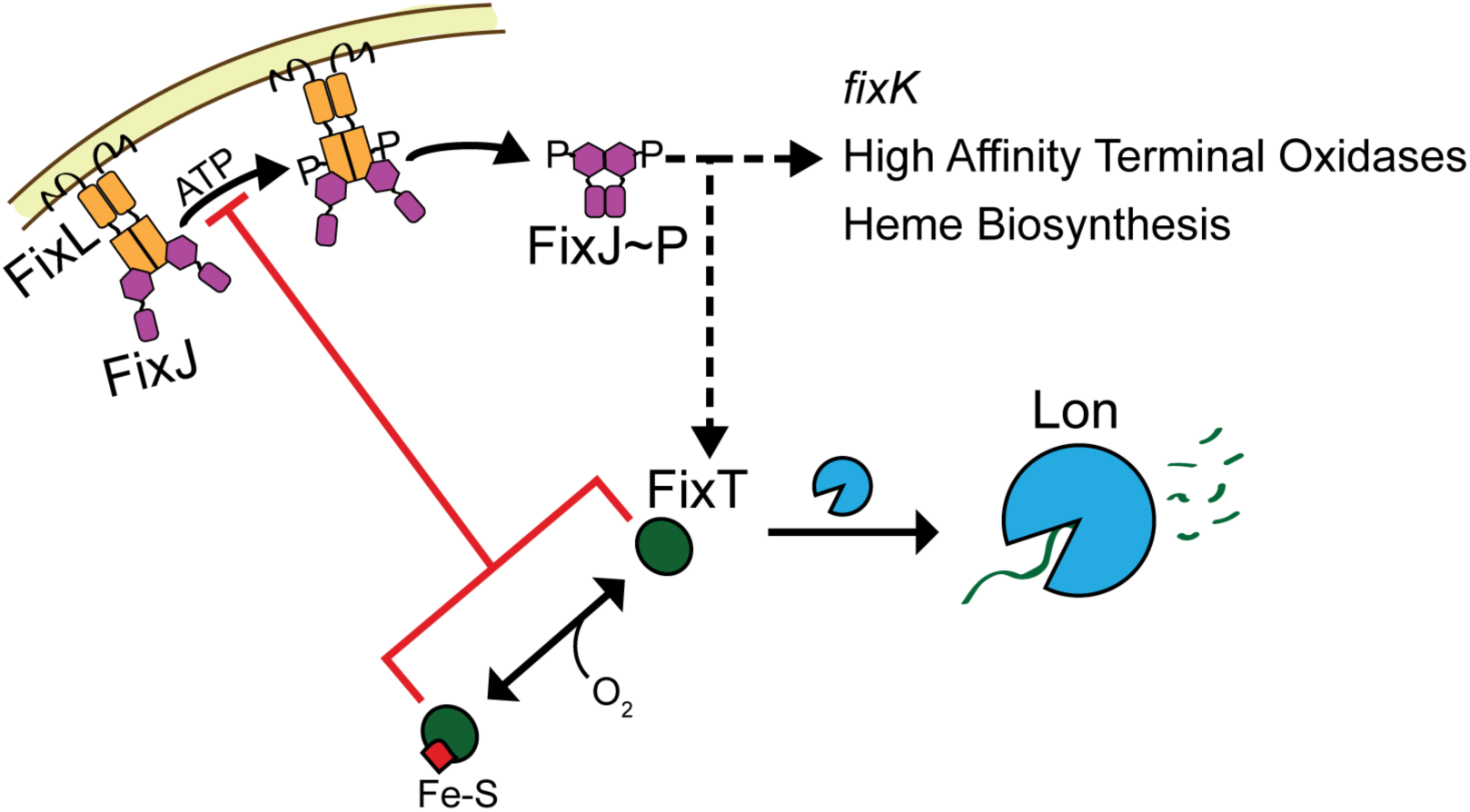
Proteolytic regulation of FixT tunes feedback inhibition of the FixLJ-mediated microoxic response. Model of FixLJ-signaling incorporating post-translational regulation of FixT. As a feedback inhibitor, FixT is capable of integrating additional information into the FixLJ-signaling system. Low oxygen levels increase FixL autokinase activity leading to increased FixJ phosphorylation and transcriptional activation (dashed-black arrow) of the microaerobic regulon, which includes the *fixK* transcription factor, high affinity terminal oxidases, heme biosynthesis genes, and *fixT*. FixT directly inhibits the autophosphorylation activity of FixL (red line). In addition, FixT can bind an Fe-S cluster that does not affect inhibition of FixL *in vitro*. Instead, our data support a model in which loss of the Fe-S cluster accelerates degradation of FixT by the Lon protease, thereby reducing FixT-dependent feedback inhibition. Cellular cues, such as oxygen (or potentially other signals), can affect the state of the FixT Fe-S cluster, and thus, FixT protein levels. In addition, Lon-dependent degradation of FixT may be sensitive to additional factors, such as adaptor proteins.

Although FixT Fe-S-binding appears restricted to *Caulobacter* species, sensory input via proteolysis may be a more universal regulatory theme. A previous study reported an intriguing genetic interaction between *fixT* and *asnO* in *S. meliloti* (23). As yet, no mechanism has been published explaining why *asnO* is required for FixT activity *in vivo*. Given our results, perhaps AsnO directly or indirectly protects FixT from proteolysis. We look forward to future studies that will test whether *S. meliloti* and other species also employ Lon (or other proteases) to regulate FixT levels, and thus FixLJ signaling output, in the cell.

## Materials and Methods

### Strains and Plasmids

All plasmids were cloned using standard molecular biology techniques. Primers are available upon request. See Table S1 for strain and plasmid information. For *in vivo* overexpression, FixT constructs were inserted into pMT805, a replicating plasmid for xylose-inducible expression (70). *ftsH* and *hslU* in-frame knockout plasmids were generated by cloning homologous upstream and downstream regions into pNTPS138. Plasmids were transformed into *C. crescentus* CB15 strains by electroporation or triparental mating. For protein expression, FixT, FixL_118-495_, and FixJ were inserted into a H_6_-Sumo pET23b expression vector. We note that the FixT sequence referenced in this work is the original ORF analyzed by Crosson et al. (2005). This sequence predicts an earlier start codon, and therefore 3 additional N-terminal amino acids (MRR), than the sequence currently annotated in NCBI. Knockout strains were generated in a Δ*fixT* CB15 background by a double recombination strategy (71). *ftsH* knockouts were recovered as described (72).

### Protein Expression and Purification

All constructs were expressed as H_6_-Sumo fusions in BL21 (DE3) cells (Rosetta or Bl21 Star) and grown in LB medium. For FixT (for aerobic purification) and FixJ expression, cultures were grown at 37°C, induced with 0.5 mM IPTG at OD_600_ = 0.5-0.6, and harvested after 1.5-3 hours. For FixT expression prior to anaerobic purification, cultures were grown at 37°C, transferred to a sealed glass bottle at OD_600_ = 0.6, and induced with 0.5 mM IPTG for 3.5 hours. To express FixL, cultures were grown at 37°C to OD_600_ = 0.6, 5-aminolevulinic acid was added to 250 µM and cells were grown for 1 additional hour before adding 0.5 mM IPTG. Cultures were then grown overnight at 18°C.

To purify FixT, cell pellets were lysed, using an LV1 microfluidizer, in TL Buffer (20 mM Tris pH 8, 150 mM NaCl, 10 mM Imidazole, 1 mM DTT) with the addition of 1 mM PMSF and 5 µg/mL DNAse I. Lysates were clarified and subjected to Ni-nitrilotriacetic acid (NTA) affinity chromatography (Ni-NTA Superflow Resin, Sigma Aldrich) before and after cleavage with Ulp1 protease. Samples were incubated with 10 mM EDTA for 2 hours, followed by dialysis for 2 hours at room temperature into T Buffer (20 mM Tris pH 8, 150 mM NaCl, 1 mM DTT). Protein was further purified by gel filtration chromatography (HiLoad 16/600 Superdex 200 pg) in T buffer. FixL was purified in a similar manner as FixT, but pellets were lysed in LL Buffer (20 mM Tris pH 8, 150 mM NaCl, 10 mM Imidazole, 10% glycerol) with the addition of 1 mM PMSF and 5 µg/mL DNAse I. Lysates were purified twice by Ni-NTA, followed by gel filtration chromatography in L Buffer (20 mM Tris pH 8, 150 mM NaCl, 10% glycerol). FixJ was purified by double Ni-NTA chromatography in the same manner as FixL and stored in L Buffer. All purified proteins were flash frozen in liquid nitrogen and stored at −80°C.

For anaerobic FixT purifications, pellets were lysed in an anaerobic chamber (Coy) by sonication. Ni-NTA purification was carried out as described above but under anaerobic conditions and without Ulp1 cleavage (proteins retained their H_6_-Sumo tag). Absorption spectra were collected as described for reconstituted FixT samples.

### Phenotypic analysis and β-galactosidase assays

Strains were grown overnight in 2 mL PYE + 0.15% xylose (chloramphenicol and oxytetracycline added to 1 µg/mL when appropriate) to saturation at 30°C. Cultures were then diluted to OD_660_ = 0.025 in 10 mL PYE + 0.15% xylose and grown at 30°C in 20 mm tubes. Growth curves were taken with this dilution as time 0. Phenotypic analyses and β-galactosidase assays were carried out after 24 hours growth.

For phenotypic analyses, equal numbers of cells (2.25 mL•OD_660_) were centrifuged, lysed in 300 µL Bugbuster Master Mix (Millipore), and clarified before analyzing supernatants by absorption and fluorescence spectroscopy in a Tecan 20M plate reader. β-galactosidase assays were performed as described after diluting cultures 1:10 in PYE (73).

### Phosphorylation assays with FixL

All phosphorylation assays were carried out at room temperature in Kinase Buffer (25 mM Tris pH 7.6, 1 mM MgCl_2_, 1 mM MnCl_2_, 1 mM CaCl_2_, 50 mM KCl). Assays were performed aerobically using the met form of FixL_118-495_ except for those using Fe-S reconstituted FixT (rec-FixT) (performed anaerobically with met FixL_118-495_). For autophosphorylation, FixL (5 µM), FixJ(D55A) (20 µM), and FixT (80 µM) were incubated 5 min before addition of ATP (1 mM cold ATP, 30 nM [γ-^32^P]ATP). Samples were quenched with an equal volume of 5X SDS-loading buffer (250 mM Tris pH 6.8, 50% Glycerol, 10% SDS, 10 mM DTT, 0.02% Bromophenol Blue), electrophoresed on Mini-Protean TGX Stain-Free 12% gels (Bio-Rad), and exposed to a phosphor screen for 1.5-3 hours before imaging with a Bio-Rad PMI System.

To assay the effect of increasing FixJ(D55A), FixL (5 µM) was incubated with or without FixT (80 µM) and varying FixJ(D55A) concentrations. Autophosphorylation was carried out as described above and 15 min time points were removed. After imaging, bands were quantified using ImageJ and normalized to the reaction without FixT and FixJ. Assays with varying FixT were carried out similarly in the presence of 5 µM FixL and 20 µM FixJ(D55A).

To observe loss of phosphorylated FixL, FixL (10 µM) was allowed to autophosphorylate, in the presence of 40 µM FixJ D55A, for 30 min before removal of nucleotides using Zeba Spin Desalting Columns (ThermoFisher). FixL was then added 1:1 to mixes with/without 160 µM FixT and/or 1 mM ADP.

For phosphotransfer assays, FixL (10 µM) was allowed to autophosphorylate for 30 min before mixing 1:1 with 5 µM FixJ and/or 160 µM FixT. Samples were taken after 30 seconds and analyzed as described above.

### Reconstitution of the FixT Fe-S cluster

40 µL aliquots of FixT were degassed under vacuum for 2 hours and brought into an anaerobic chamber (Coy) at room temperature. Samples were reduced for 15 min with 10 mM DTT and brought to 30-40 µM in R Buffer (50 mM Tris pH 8.3, 150 mM NaCl, 5 mM DTT). FeCl_3_ was added to 5-7-fold excess of FixT and allowed to incubate for 10 min. Na_2_S was then added equimolar to FeCl_3_ and the reaction was allowed to proceed for 1.5-2 hours. For non-reconstituted FixT controls, FeCl_3_ and Na_2_S were omitted from the reactions. Reactions were then desalted using PD-10 desalting columns (GE Healthcare) and concentrated using centrifugal filters. Following concentration, samples were removed from the anaerobic chamber and analyzed by Pierce Coomassie Plus Bradford reagent (ThermoFisher) and Ferrozine (ThermoFisher) assay as described (74). For absorption spectroscopy, proteins were diluted to 50 µM in 96-well clear bottom plates (Thermo Scientific), covered with crystal clear sealing film (Hampton Research), removed from the anaerobic chamber, and immediately analyzed in a Tecan 20M plate reader. The sealing film was then removed, and decay spectra were recorded. Buffer-only spectra were also recorded and subtracted as background.

### Western Blotting and *in vivo* protein degradation experiments

For steady-state overexpression experiments, cultures were grown as described for growth curves and β-galactosidase assays. Cells were collected (1.5 mL•OD_660_), spun for 2 min at 13000 × *g*, and lysed in 200 µL Bugbuster Master Mix + 2mM PMSF. Lysates were cleared and denatured in 1X SDS Loading Buffer. To assay native steady-state levels, primary overnight cultures were diluted to OD_660_ = 0.025 in 100 mL of PYE + 0.15% xylose in a 250 mL flask and grown for 24 hours. Samples were spun down and cell pellets were resuspended in T Buffer (2.5 mL/OD_660_). 3 mL of each sample were lysed using an LV1 microfluidizer, cleared by centrifugation, and denatured with 1X SDS Loading Buffer. For *in vivo* degradation experiments, overnight cultures were diluted to OD_660_ = 0.1 in 10 or 4 mL (20 mm tubes) and grown to OD_660_ ∼ 0.4. The cells were then treated with 25 µg/mL oxytetracycline to stop translation. Samples were removed at indicated times after oxytetracycline addition (0.3 mL•OD_660_), spun for 2 min at 13000 × *g*, resuspended with 40 µL 2.5X SDS Loading Buffer, and frozen at −80°C. After thawing, samples were treated with 250U Benzonase (Sigma Aldrich) for 15 min before boiling 3 min at 95°C.

All western blot samples were electrophoresed on Mini-PROTEAN TGX Stain-free 4-20% (w/v) pre-cast gels (Bio-Rad). Proteins were transferred to Immobilon-P membranes (EMD Millipore) or Immun-Blot LF PVDF (Bio-Rad) in a wet transfer apparatus. Membranes were probed with polyclonal antibodies (produced by Josman, LLC) raised against FixT (1:4000 dilution), incubated with either goat anti-rabbit IgG-HRP conjugate (ThermoFisher; 1:5000 dilution) or goat anti-rabbit IgG Cross-Adsorbed Secondary Antibody, Alexa Fluor 488 (Invitrogen; 1:1000 dilution), and, when appropriate, developed with SuperSignal West Femto Maximum Sensitivity Substrate (ThermoFisher). The blots were imaged using a Bio-Rad Chemidoc MP Imager. Bands were quantified using ImageJ software and normalized to values at time 0 for each blot. For each *lon*^*+*^ degradation experiment, curves were fit to a single exponential decay (R^2^ > 0.95) in Prism (Graphpad) to determine a half-life, which was then averaged across three independent experiments.

## Acknowledgements

We thank Tao Pan and Eric L. Hegg for use of equipment, and John Anderson, Peter Chien, and members of the Crosson laboratory for helpful discussion.

Research reported in this publication was supported by the National Institute of General Medical Sciences of the National Institutes of Health (NIH) under Award Numbers F32 GM128283 (B.J.S.) and R01 GM087353 and R35 GM131762 (S.C.). The content is solely the responsibility of the authors and does not necessarily represent the official views of the National Institutes of Health.

The authors declare no conflict of interest.

