## Supplementary Figures and Table for "Feedback control of a two-component signaling system by an Fe-S-binding receiver domain"

### Supplementary Figures and Tables

#### Figure S1

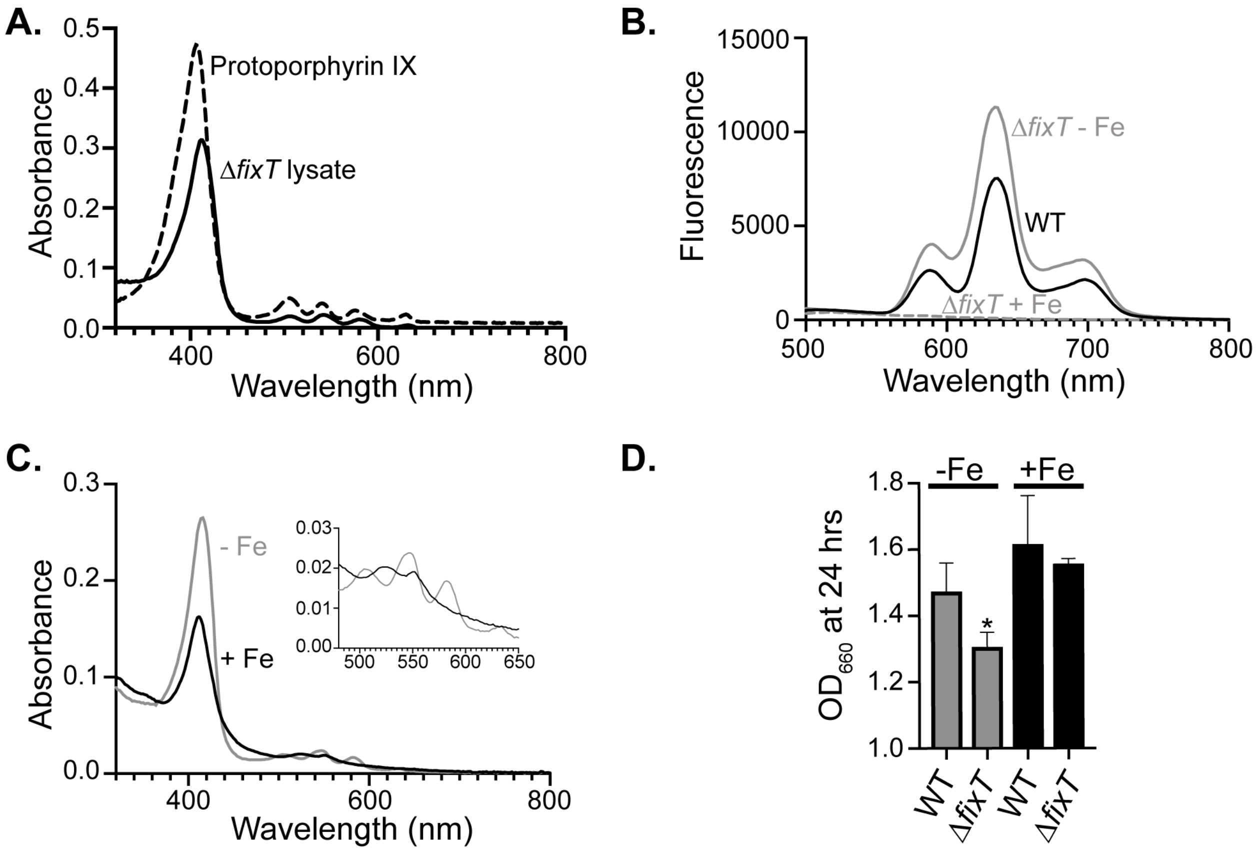


**Figure S1: Comparison of porphyrin signatures to protoporphyrin IX and effects of Fe supplementation**

(A) Absorption spectrum of Δ*fixT* lysate compared with protoporphyrin IX (Sigma Aldrich) in DMSO.

(B) Fluorescence emission scan of wild-type (WT), Δ*fixT*, and Fe-treated Δ*fixT* lysates (Ex. 404 nm).

(C) Absorption spectra of lysates from Δ*fixT* cells grown with (black) or without (grey) 10 μM $FeSO_4$. Inset highlights *Q* bands in 500-650 nm region.

(D) Optical density ($OD_{660}$) of cultures grown for 24 hours with (black bars) or without (grey bars) addition of 10 μM $FeSO_4$ (average ± SD, n > 3). * = $p < 0.001$, one-way ANOVA; $p < 0.05$, Dunnett's post-test comparison to wild-type (WT) untreated.

### Figure S2

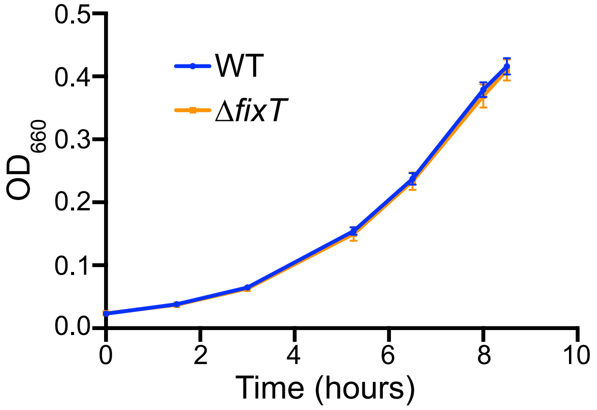


**Figure S2: Growth of wild-type and Δ*fixT* strains**

Growth curves during exponential phase of wild-type (WT, blue) and Δ*fixT* (orange) strains measured by optical density ($OD_{660}$). Points are averages of 3 biological replicates ± SD. Strains were grown under oxygen-limited conditions as in Fig. 1.

### Table S1: Strains and plasmids used in this work

| ***E. coli* strains with plasmids** | | | |
|---|---|---|---|
| **Strain #** | **Genotype** | **Comments** | **Source** |
| FC929 | TOP10 | Cloning strain | Invitrogen |
| FC3 | MT607 / pRK600 | Helper strain for tri-parental matings | (1) |
| FC803 | Rosetta (DE3) pLysS | Strain for protein expression from T7 promoter | Novagen |
| ***Caulobacter* expression plasmids** | | | |
| MTLS4423 | TOP10 / pMT805 (pBXMCS-2) | Replicating plasmid (mid copy, BBR origin) with xylose inducible promoter, $Chlor^R$ | (2) |
| FC3396 | TOP10 / pMT805-*fixT* | Cloning Primers:<br>5'-gttgcatATGCGACGCATGTGCGAC-3'<br>5'-caacgagctcctaGAGCCCCGGCG-3' | This work |
| FC3397 | TOP10 / pMT805-*fixT(C53S)* | Generated from FC3396 | This work |
| FC3398 | TOP10 / pMT805-*fixT(C56S)* | Generated from FC3396 | This work |
| FC3399 | TOP10 / pMT805-*fixT(C59S)* | Generated from FC3396 | This work |
| FC3400 | TOP10 / pMT805-*fixT(C64S)* | Generated from FC3396 | This work |
| FC3401 | TOP10 / pMT805-*fixT(4C)* | Mutant containing all mutations in above 4 plasmids, Generated from FC3396 | This work |
| **Allele Replacement Plasmids** | | | |
| FC55 | DH10B / pNPTS138 | Allele replacement plasmid, $Kan^R$, SacB | M.R.K. Alley |
| FC106 | TOP10/ pNPTS138-Δ*fixT* | | (3) |
| FC3402 | TOP10 / pNPTS138-Δ*hslU* | knockout allele contains first 6 and last 3 codons of *hslU*. 5' end (5'-ATGACCGAG…), 3' end (…ATTTTGtag-3') | This work |
| FC3403 | TOP10 / pNPTS138-Δ*ftsH* | knockout allele contains first 4 and last 107 codons of *ftsH*. 5' end (5'-ATGAATTTC…), 3' end (…ACCGCCtga-3') | This work |
| FC2256 | TOP10 / pNPTS138-Δ*clpA* | knockout allele contains first 13 and last 11 codons of *clpA*. 5' end (5'-TTGCCCTCT…), 3' end (…GCCGAAtag-3') | This work |
| **Transcriptional *lacZ* fusion plasmids** | | | |
| FC94 | TOP10 / pRKlac290-P*fixK* | Plasmid containing $P_{fixK}$-*lacZ* transcriptional fusion, $tet^r$ | (3) |
| **Heterologous protein expression plasmids** | | | |
| FC3404 | DH5α / pET23b-$H_6$-*smt3* | Plasmid to express proteins fused to the $H_6$-SUMO tag | (4) |
| FC3249 | DH5α / pET23b-$H_6$-*smt3* V2 | Derived from FC3404, contains additional restriction sites for cloning | Breann Brown |
| FC3405 | DH5α / pET23b-Ulp1 | Plasmid to express $H_6$-Ulp1 for cleavage of SUMO tag | (4) |
| FC3406 | TOP10 / pET23b-$H_6$-*smt3-fixT* | *fixT* insert amplified from FC3396 | This work |
| FC3407 | TOP10 / pET23b-$H_6$-*smt3-fixT(C64S)* | *fixT* insert amplified from FC3400 | This work |
| FC3408 | TOP10 / pET23b-$H_6$-*smt3-fixT(4C)* | *fixT* insert amplified from FC3401 | This work |
| FC3409 | TOP10 / pET23b-$H_6$-*smt3-*$fixL_{118\text{-}495}$ | Cloning Primers:<br>5'-gatcaccggtggtGCGGCGGCGGTCAATG-3'<br>5'-caatgagctctcaGTCATCGATGGTCTCC-3' | This work |
| FC3410 | TOP10 / pET23b-$H_6$-*smt3-fixJ* | Cloning Primers:<br>5'-gatcaccggtggtATGACTGACGCCCC-3'<br>5'-gtgcggccgctcaGCCGCCGCGC-3' | This work |
| FC3411 | TOP10 / pET23b-$H_6$-*smt3-fixJ(D55A)* | Generated from FC3410 | This work |

| ***Caulobacter crescentus* strains** | | | |
|---|---|---|---|
| **Strain #** | **Genotype** | **Source** | **Figure** |
| FC19 | Wild type CB15 | (5) | 1, 5, S1,S2 |
| FC3412 | CB15 / pMT805 | This work | 1, 4 |
| FC98 | CB15 / pRKlac290-P*fixK* | (3) | 1 |
| FC3413 | CB15 / pMT805 / pRKlac290-P*fixK* | This work | 4 |
| FC13 | CB15 Δ*fixT* | (3) | 1, S1, S2 |
| FC3414 | CB15 Δ*fixT* / pMT805 | This work | 1, 5 |
| FC112 | CB15 Δ*fixT* / pRKlac290-P*fixK* | (3) | 1 |
| FC3415 | CB15 Δ*fixT* / pMT805 / pRKlac290-P*fixK* | This work | 4 |
| FC3416 | CB15 Δ*fixT* / pMT805-*fixT* | This work | 1, 5 |
| FC3417 | CB15 Δ*fixT* / pMT805-*fixT(4C)* | This work | 5 |
| FC3418 | CB15 Δ*fixT* / pMT805-*fixT* / pRKlac290-P*fixK* | This work | 4 |
| FC3419 | CB15 Δ*fixT* / pMT805-*fixT(C53S)* / pRKlac290-P*fixK* | This work | 4 |

| | | | |
|---|---|---|---|
| FC3420 | CB15 Δ*fixT* / pMT805-*fixT(C56S)* / pRKlac290-P*fixK* | This work | 4 |
| FC3421 | CB15 Δ*fixT* / pMT805-*fixT(C59S)* / pRKlac290-P*fixK* | This work | 4 |
| FC3422 | CB15 Δ*fixT* / pMT805-*fixT(C64S)* / pRKlac290-P*fixK* | This work | 4 |
| FC3423 | CB15 Δ*fixT* / pMT805-*fixT(4C)* / pRKlac290-P*fixK* | This work | 4 |
| FC2264 | CB15 Δ*lon* | (6) | 5 |
| FC3424 | CB15 Δ*fixT*Δ*lon* | This work | 5 |
| FC3425 | CB15 Δ*fixT*Δ*lon* / pMT805-*fixT* | This work | 5 |
| FC3426 | CB15 Δ*fixT*Δ*lon* / pMT805-*fixT(4C)* | This work | 5 |
| FC3427 | CB15 Δ*fixT*Δ*hslU* | This work | |
| FC3428 | CB15 Δ*fixT*Δ*hslU* / pMT805-*fixT* | This work | 5 |
| FC2265 | CB15 Δ*clpA* | This work | |
| FC3429 | CB15 Δ*fixT*Δ*clpA* | This work | |
| FC3430 | CB15 Δ*fixT*Δ*clpA* pMT805-*fixT* | This work | 5 |
| FC2417 | CB15 Δ*socAB*Δ*clpX* / pAC152 | (7) | |
| FC3431 | CB15 Δ*fixT*Δ*socAB*Δ*clpX* / pAC152 | This work | |
| FC3432 | CB15 Δ*fixT*Δ*socAB*Δ*clpX* / pAC152 / pMT805-*fixT* | This work | 5 |
| FC3433 | CB15 Δ*fixT*Δ*ftsH* | This work | |
| FC3434 | CB15 Δ*fixT*Δ*ftsH* / pMT805-*fixT* | This work | 5 |
